# Visuo-proprioceptive recalibration and the sensorimotor map

**DOI:** 10.1101/2022.11.29.517247

**Authors:** Hannah J. Block, Yang Liu

## Abstract

Spatial perception of our hand is closely linked to our ability to move the hand accurately. We might therefore expect that reach planning would take into account any changes in perceived hand position; in other words, that perception and action relating to the hand should depend on a common sensorimotor map. However, there is evidence to suggest that changes in perceived hand position affect a body representation that functions separately from the body representation used to control movement. Here we examined target-directed reaching before and after participants either did (Mismatch group) or did not (Veridical group) experience a cue conflict known to elicit recalibration in perceived hand position. For the reaching task, participants grasped a robotic manipulandum that positioned their unseen hand for each trial. Participants then briskly moved the handle straight ahead to a visual target, receiving no performance feedback. For the perceptual calibration task, participants estimated the locations of visual, proprioceptive, or combined cues about their unseen hand. The Mismatch group experienced a gradual 70 mm forward mismatch between visual and proprioceptive cues, resulting in forward proprioceptive recalibration. Participants made significantly shorter reaches after this manipulation, consistent with feeling their hand to be further forward than it was, but reaching performance returned to baseline levels after only 10 reaches. The Veridical group, after exposure to veridically-aligned visual and proprioceptive cues about the hand, showed no change in reach distance. These results are not fully consistent with a single common sensorimotor map, but could suggest multiple, interacting body representations.

**NEW & NOTEWORTHY:** If perceived hand position changes, we might assume this affects the sensorimotor map and, in turn, reaches made with that hand. However, there is evidence for separate body representations involved in perception vs. action. After a cross-sensory conflict that results in proprioceptive recalibration in the forward direction, participants made shorter reaches as predicted, but only briefly. This is not fully consistent with a single common sensorimotor map, but could suggest multiple, interacting body representations.

## 1. INTRODUCTION

To plan and execute hand movements to interact efficiently with objects in the environment, the brain must have an accurate representation of the hand’s position. This representation is thought to be multisensory, including both visual information from the eyes and proprioceptive information from the muscles and joints of the upper limb (Ghahramani et al. 1997). When both a visual estimate (*h*_*V*_) and a proprioceptive estimate (*h*_*P*_) of true hand position (*H*) are available, these are weighted and combined to form a single integrated estimate (*h*_*VP*_):

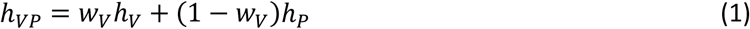

where *w*_*V*_ is the weight of vision relative to proprioception (i.e., *w*_*V*_ = 0.7 implies 70% reliance on vision and 30% reliance on proprioception). Weighting may be determined by relative variance in the sensory signals (van Beers et al. 1999; Ernst and Banks 2002; Ghahramani et al. 1997) as well as top-down influences such as attention or task demands (Limanowski 2022; Limanowski and Friston 2020).

*h*_*V*_ and *h*_*P*_ do not agree perfectly even in normal circumstances (Smeets et al. 2006), but in the presence of an externally-imposed conflict between these cues, visuo-proprioceptive recalibration occurs (Hsiao et al. 2022; Salomonczyk et al. 2013; Welch 1978). This may involve both a shift of the proprioceptive estimate closer to the visual estimate (*Δh*_*P*_), and vice versa (*Δh*_*V*_).

We might expect that reach planning would take into account any such changes in perceived hand position; in other words, that perception and action relating to the hand should depend on a common sensorimotor map. We recently observed somatotopically-focal changes in the excitability of primary motor cortex (M1) that were specifically related to visuo-proprioceptive recalibration, after controlling for motor behavior (Mirdamadi et al. 2022; Munoz-Rubke et al. 2017); findings like these are suggestive of a close relationship between perception of hand position and motor execution involving that hand. However, substantial work has suggested that changes in perceived hand position may affect a body representation that functions separately from the body representation used to control movement; these have been referred to as the body image and body schema, respectively (Kammers et al. 2009; Paillard 1999; de Vignemont 2010).

Literature addressing some form of visuo-proprioceptive recalibration is mixed as to whether movement is affected. One challenge is the lack of research specifically focused on visuo-proprioceptive recalibration, unconfounded by other processes that are often the primary focus. For example, there is relevant literature using the Rubber Hand Illusion (RHI), a paradigm that creates illusory body ownership over a fake arm through synchronous stroking of the seen fake arm and the hidden real arm (Botvinick and Cohen 1998). The RHI in a sense involves a visuo-proprioceptive cue conflict, since the fake arm is meant to be a visual cue related to the real arm, and proprioceptive recalibration (described as drift in this literature) is thought to occur (Abdulkarim et al. 2021; Abdulkarim and Ehrsson 2018). Kammers et al. (2009) concluded that the RHI affected the body image but not the body schema, after observing that perceptual judgments, but not ballistic motor responses, were sensitive to the RHI. On the other hand, in the original RHI study by Botvinick and Cohen (1998), the illusion was assessed by having participants point at the perceived location of their other hand, a motor response that clearly *was* affected by the RHI (Botvinick and Cohen 1998; Kammers et al. 2009). A complication is that in both studies, the pointing movements (Botvinick and Cohen 1998) and ballistic movements (Kammers et al. 2009) were directed toward the other hand, potentially causing participants to access both body schema and body image.

A second relevant body of literature is the subset of visuomotor adaptation research that measures proprioceptive recalibration. Visuomotor adaptation is a cerebellum-dependent process in which participants experience a systematic perturbation of their movements and gradually compensate by updating their sensorimotor map to reduce systematic errors (Robertson and Miall 1999; Seidler and Noll 2008). A common method of perturbation is cursor rotation, where participants move their unseen hand to guide a cursor to a target on a screen. The cursor can be rotated; e.g., a movement straight ahead results in the cursor moving 30° to the right. In addition to causing movement errors that lead to updating of motor commands, the mismatch between hand and cursor creates a visuo-proprioceptive conflict. Indeed, proprioceptive recalibration has now been documented extensively in this paradigm (Ostry and Gribble 2016; Rossi et al. 2021; Salomonczyk et al. 2013).

Several studies have used a modified cursor rotation paradigm to eliminate the motor adaptation aspect: participants move their hand along a set channel that gradually deviates from the cursor, which always moves to the target so that no movement error is apparent but the visuo-proprioceptive mismatch is still created (Cressman and Henriques 2010; Mostafa et al. 2019; Ruttle et al. 2018).

Participants indeed recalibrate proprioception in these circumstances, and additionally, target-directed reaches with no cursor show a shift in direction consistent with the change in felt hand movement direction (Cressman and Henriques 2010; Mostafa et al. 2019; Ruttle et al. 2018).

It makes sense that a visuo-proprioceptive mismatch created in the context of cursor rotation, with the hand actively or passively moved in a direction rotated from the cursor and target, would alter the same sensorimotor map used to plan target-directed reaches. However, it is difficult to generalize this finding to visuo-proprioceptive recalibration in general, which does not require either active or passive movement, or a movement target. In the cursor rotation paradigm, the cue conflict exists only in the context of target-directed movement: the conflict changes from zero at the home position to some maximum angular deviation at the target position.

Here we ask a slightly different question: Does visuo-proprioceptive recalibration, triggered with a cue conflict at a static position, affect the same sensorimotor map used to make active target-directed reaching movements? Or does it affect a separate body representation, as in the body image vs. body schema concept? To answer this question, two groups of participants made straight-ahead target-directed reaches, with no cursor, before and after experiencing either a visuo-proprioceptive conflict (Mismatch group) or veridical visuo-proprioceptive cues (Veridical group) while the hand was stationary. The cue conflict was introduced by gradually shifting the visual cue forward from the hand, to a maximum of 70 mm (Fig. 1). Proprioceptive recalibration was thus expected in the forward direction (Fig. 1Bii). We therefore predicted that the Mismatch group, feeling the hand to be further forward than it really was after proprioceptive recalibration, would make shorter reaches after experiencing the conflict (Fig. 1Ci-ii), while Veridical group reach distance would remain unchanged.

**Figure 1.**
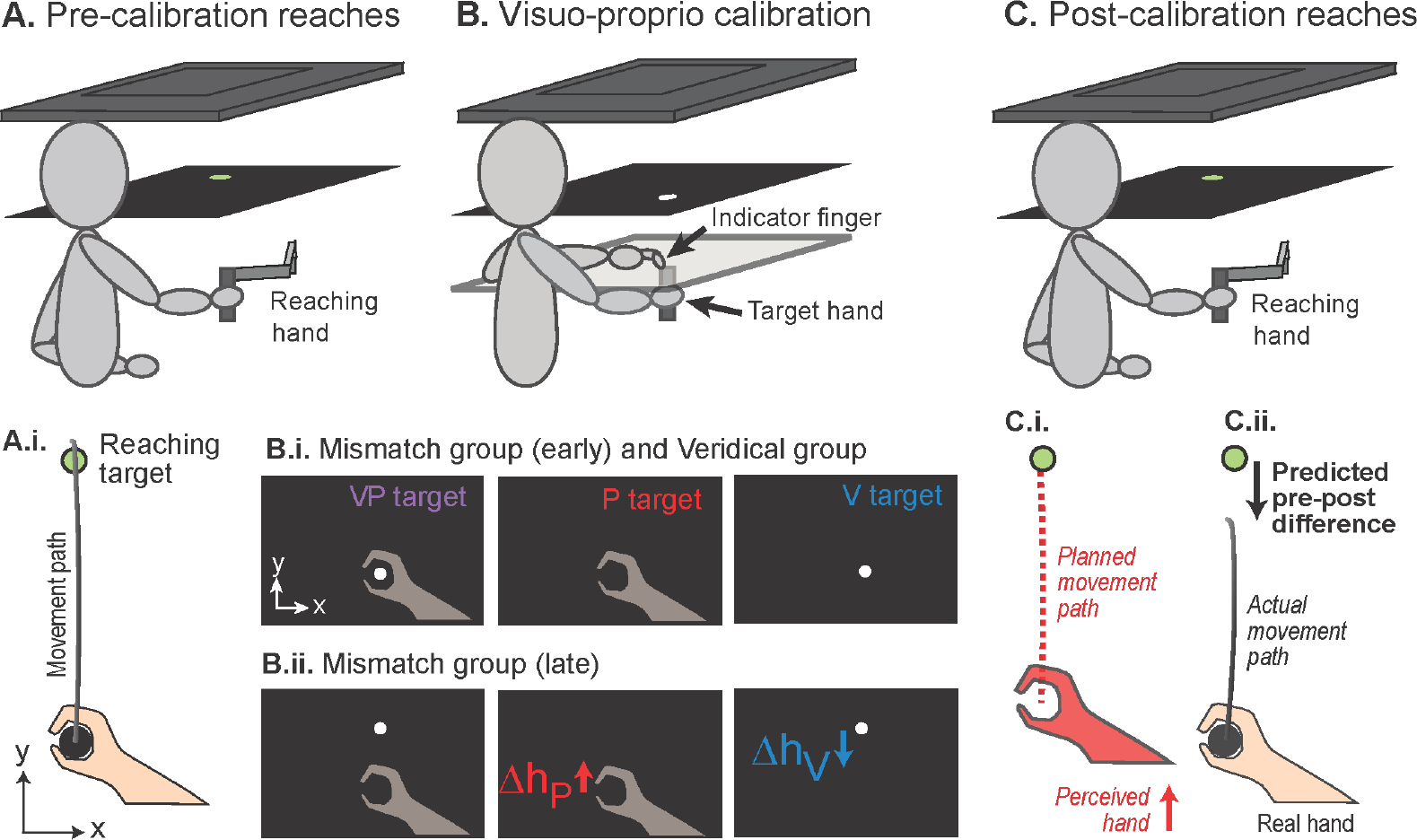
Top row: Participant holds handle of robotic manipulandum in right hand. Images viewed in mirror of 2D VR display appear to be in line with top of manipulandum handle. Not pictured: fabric preventing vision of the upper arms. No direct vision of hands is possible. **A. Pre-calibration reaches**. Participant performs straight-ahead reaching with no visual feedback about hand position, stopping at perceived position of visual target. The hand was brought passively back to the invisible starting position after each trial. **B. Visuo-proprioceptive calibration. i**. Participants use their left index finger on a touchscreen to indicate the perceived position of visual (V), proprioceptive (P), and combined (VP) targets. No performance feedback or knowledge of results was available at any time. For the Veridical group, VP targets remained veridical throughout. **ii**. For the Mismatch group, the visual component (white disc) gradually shifted forward to a maximum of 70 mm. This generally results in both proprioceptive recalibration toward the visual target (*Δh*_*P*_) and the visual recalibration toward the proprioceptive target (*Δh*_*V*_). Transparent hands and writing were not visible to participants. **C. Post-calibration reaches**. Same procedure as pre-calibration reaches. **i-ii**. For the Mismatch group, if *Δh*_*P*_ affects reaching, we predict participants will stop short of the visual target because they feel their hand is further from them (closer to the target) than it is. With only proprioceptive information about hand position, planned movements (red dashed lines) should be shorter relative to pre-perturbation (black arrow) for the Mismatch group but not the Veridical group.

## 2. METHODS

### Participants

32 healthy right-handed adults participated in the study, which consisted of one lab visit. Inclusion criteria were: aged 18-45 and right handed with normal or corrected-to-normal vision. Exclusion criteria were any muscular, orthopedic, or neurological disorders. All enrolled participants reported that they met these inclusion and exclusion criteria. The study was approved by Indiana University Institutional Review Board, and all participants gave written informed consent. Participants were randomly assigned to the Mismatch group (N = 16, mean age 19.9 ± SD 1.2 years, 4 males) or the Veridical group (N = 16, mean age 20.5 ± 1.8 years, 6 males) using a random sequence of ones and twos (16 of each) generated in MATLAB 2021a (Mathworks). The study was single blind, with the experimenter knowing the participant’s group assignment but the participant not knowing.

### Apparatus

Participants were seated at a reflected rear projection apparatus to perform a task with three parts (Fig. 1), grasping the handle of a KINARM Endpoint 2D robotic manipulandum (BKIN) with their right hand throughout. Positional accuracy of the manipulandum, with high-resolution secondary encoders, is 3 microns; inertial load of the passive manipulandum is 0.8/1.0 kg (minor/major axes). Participants had no direct vision of their hand, but viewed a task display that appeared to be in the plane of the manipulandum (Fig. 1A).

### Procedures

The experimental session consisted of three parts. First, pre-calibration straight-ahead right-hand reaches to a visual target with no visual feedback about the right hand (Fig. 1A). Second, a visuo-proprioceptive calibration task with either mismatched or veridical visual and proprioceptive cues about the right hand, depending on group assignment (Fig. 1B). Third, post-calibration straight-ahead right-handed reaches to a visual target with no visual feedback about the right hand (Fig. 1C). The experiment was preceded by instructions about the two tasks (reaching and calibration) and practice of each task. The whole session took about one hour.

#### Straight-ahead reaching

Before and after the visuo-proprioceptive calibration task (Fig. 1A and C), participants were asked to grasp the manipulandum handle in their right hand and briskly move from the starting position to the target. The hand was brought passively to the starting position for each trial, as the starting position was not visible. No online or endpoint visual feedback about hand position was given at any point in this task. Reach endpoint was defined as the position at which movement velocity dropped below 5% of peak velocity.

In addition to practice trials, participants performed 20 reaches pre-calibration and 20 reaches post-calibration. The starting position was located at the participant’s body midline, about 20 cm in front of their chest. The visual target was located 10 cm forward of the starting position.

Reaches were binned into sets of 5. To determine if reach endpoints were closer or further from the participant after the calibration task, the y-coordinates of each set of 5 reach endpoints were averaged within participants.

#### Visuo-proprioceptive calibration tas

Visuo-proprioceptive estimates of hand position are most commonly measured and/or perturbed with a bimanual task, using an “indicator” (left) hand to indicate the participant’s perception of the “target” (right) hand’s position when visual, proprioceptive, or both types of information about the target are available (van Beers et al. 1996, 1998, 1999, 2002; Smeets et al. 2006). Participants were therefore asked to use their *unseen left index finger* to indicate on a 32 inch touchscreen (PQLabs) where they perceived a series of targets (Table 1) related to the *right (target) hand* (Fig. 1B), which grasped a stationary manipulandum handle at the target position beneath the touchscreen. It was not physically possible to place the touchscreen in the horizontal plane of the visual task display at the top of the manipulandum handle, due to the design of the manipulandum. Therefore for this task we moved the actual manipulandum out of the workspace and placed a replica manipulandum handle under the touchscreen for participants to grasp on P and VP targets. The replica was the same height and diameter as the real handle, and covered with an identical rubber grip.

**Table 1.**
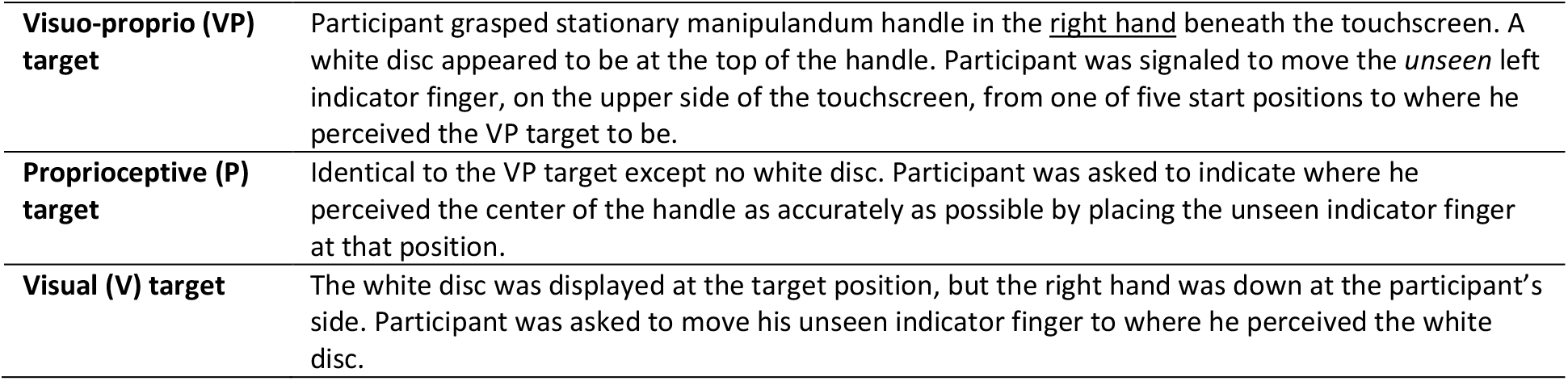
Targets in visuo-proprioceptive calibration task.

|  |  |
| --- | --- |
| **Visuo-proprio (VP) target** | Participant grasped stationary manipulandum handle in the <u>right hand</u> beneath the touchscreen. A white disc appeared to be at the top of the handle. Participant was signaled to move the *unseen* left indicator finger, on the upper side of the touchscreen, from one of five start positions to where he perceived the VP target to be. |
| **Proprioceptive (P) target** | Identical to the VP target except no white disc. Participant was asked to indicate where he perceived the center of the handle as accurately as possible by placing the unseen indicator finger at that position. |
| **Visual (V) target** | The white disc was displayed at the target position, but the right hand was down at the participant's side. Participant was asked to move his unseen indicator finger to where he perceived the white disc. |

Both groups performed 84 trials: 42 VP, 21 V, and 21 P, in repeating order (VP-V-VP-P). For the Veridical group, the white disc was always displayed veridically at the top of the replica manipulandum handle. For the Mismatch group, the white disc moved 1.67mm forward on each VP trial. Participants do not generally notice this perturbation, which results in a 70mm visuo-proprioceptive mismatch by the end of the 84 trials (Fig. 1Bii) (Block et al. 2013; Block and Bastian 2011, 2012; Hsiao et al. 2022). Indicator finger start position was jittered to prevent participants from being influenced by indicator finger movements on the previous trial. Importantly, there was no speed requirement and no performance feedback or knowledge of results, to preclude motor adaptation.

If the proprioceptive estimate of the right hand, as shown by left indicator finger endpoints, moves forward to close the visuo-proprioceptive gap (*Δh*_*P*_), then we observe overshoot on P targets. Similarly, if perceived position of the white disc moves closer to the right hand (*Δh*_*V*_), then we observe undershoot on V targets. *VP trials are used to create the mismatch while V and P trials are used to assess visual and proprioceptive recalibration*. Thus, outcome measures are based on V and P trials. We quantified visual and proprioceptive recalibration (*Δh*_*V*_ and *Δh*_*P*_) as previously (Block et al. 2013; Block and Bastian 2011, 2012): after calculating mean indicator finger endpoint positions in the y-dimension on the first and last 4 V and P trials, respectively, we computed the difference relative to actual target position, which was constant for P targets but shifts 70mm for V targets (Mismatch group only):

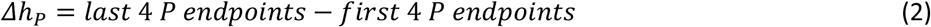

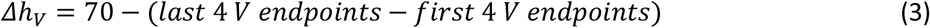

### Statistical analysis

Data were processed using MATLAB 2021a (MathWorks Inc., Natick, MA, United States). A mixed model ANOVA was performed on the straight-ahead reaching endpoints, with within-participant factor “reach set” (the last set of 5 pre-calibration reach endpoints and the four sets of 5 post-calibration reach endpoints) and between-participant factor “group” (Mismatch and Veridical). For a significant interaction, paired-sample t-tests were performed within each group, comparing the last set of pre-calibration reaches with each set of post-calibration reaches.

For the Mismatch group only, the magnitude of change in reach endpoint (first set of post-calibration reaches minus last set of pre-calibration reaches) was compared to the magnitude of proprioceptive recalibration (*Δ*h_P_) and the magnitude of total recalibration (*Δ*h_P_ + *Δ*h_V_) in a one-way repeated measures ANOVA. For a significant effect, change in reach endpoint was compared to the other two magnitudes with a paired-sample t-test.

For the post-hoc t-tests, false discovery rate was controlled by the Benjamini-Hochberg procedure (Benjamini and Hochberg 1995) with α set to 0.05. In the text, adjusted p-values are indicated as *p*_*adj*_. Data and analysis code are publicly available at https://osf.io/zy49x/.

## 3. RESULTS

On average, participants in the Mismatch group recalibrated proprioception 17.4 ± 4.0 mm and vision 38.1 ± 6.1 mm (mean ± SE) in response to a gradually-imposed 70 mm mismatch between visual and proprioceptive cues. The example participant in Fig. 2 recalibrated to a degree consistent with the group behavior. After exposure to the cue conflict, this participant reached shorter distances with the recalibrated hand, consistent with predictions (Fig. 1C); however, this was only evident in the first set of 5 trials post-mismatch (Fig. 2C).

**Figure 2.**
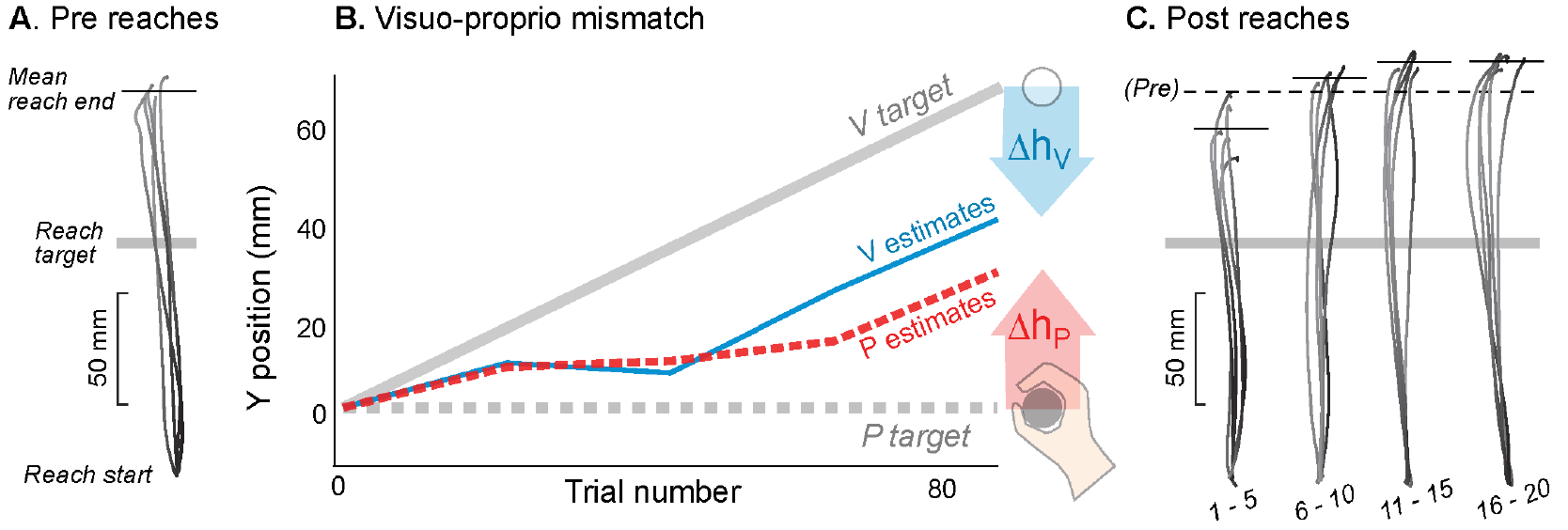
Example participant in the Mismatch group. **A**. Pre-mismatch reaches. Movement paths of the 5 right-handed reaches immediately preceding mismatch task. With no performance feedback or knowledge of results, this participant consistently overshot the reach target (grey bar). **B**. The mismatch task gradually imposed 70 mm of visuo-proprioceptive mismatch by shifting the white disc (V target) forward from the stationary right hand (P target). This participant recalibrated both proprioception (*Δh*_*P*_ = 29.6 mm) and vision (*Δh*_*V*_ = 28.9 mm). **C**. Four sets of 5 right-handed reaches following mismatch task. The first set of 5 reaches undershot the pre-mismatch mean (dashed line) by 16.4 mm.

Participants’ average reach distances on the four sets of five right-handed reaches (Fig. 3i and iii) were analyzed with a mixed-model ANOVA with factors Group (Mismatch, Veridical) and Reach Set (5 sets). There was a significant Reach Set x Group interaction (F_3,90_ = 3.34, p = 0.023, η_p_^2^ = 0.01), suggesting that the two groups differed in reach distance across the five reach sets. There was also an effect of Reach Set on reach distance (F_3,90_ = 3.98, p = 0.010, η_p_^2^ = 0.01), but no main effect of Group (F_1,90_ = 0.25, p = 0.62, η_p_^2^ = 0.008).

**Figure 3.**
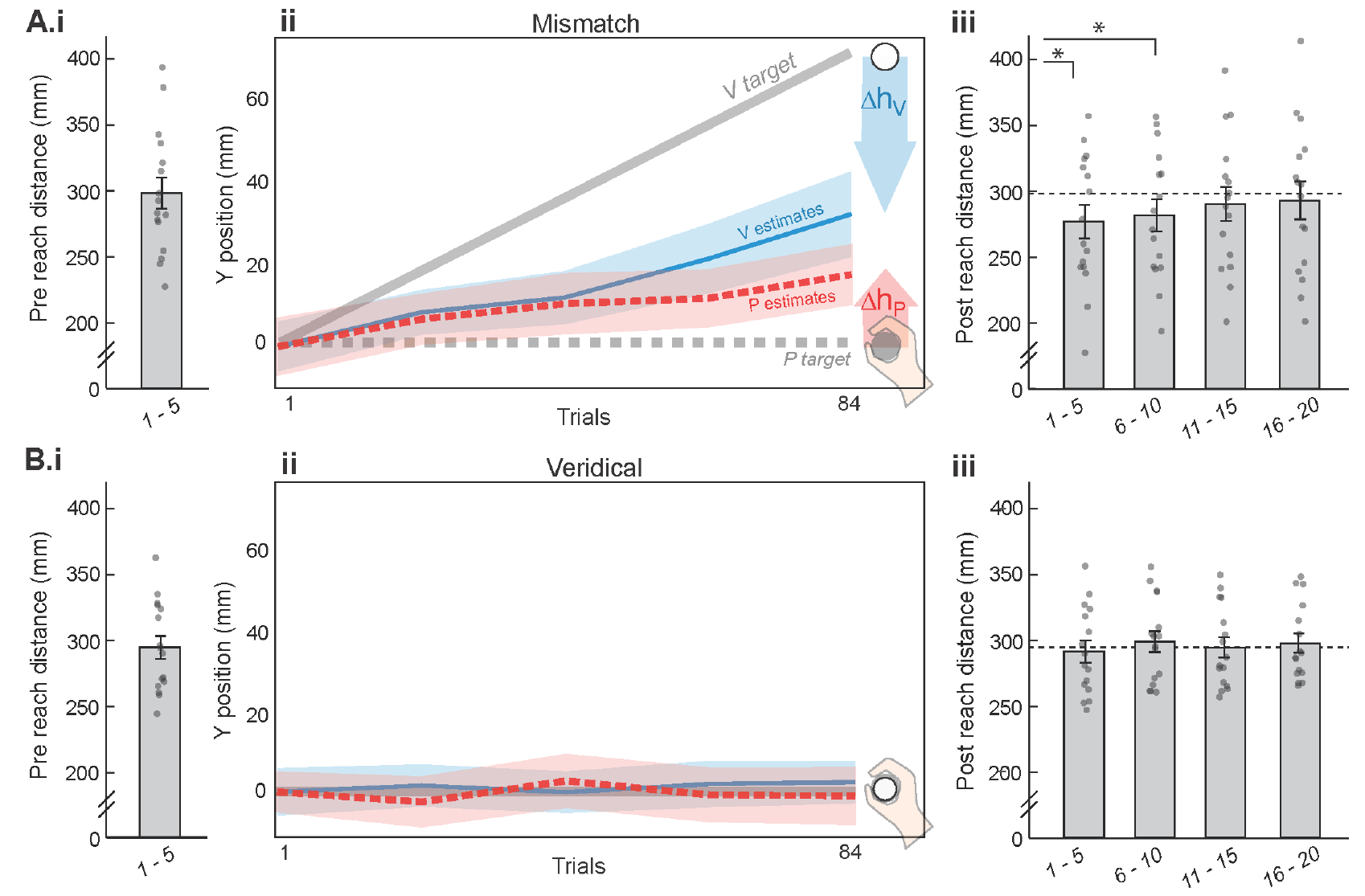
Group reaching distances pre- and post-calibration task. **A. Mismatch group (N=16). i**. Mean distance of the 5 right-handed reaches immediately preceding calibration task. Dots represent individual participants. **ii**. Mismatch task. Participants pointed with their left indicator finger to P targets (right hand, dashed grey line), V targets (white disc, solid grey line), and VP targets (combined), with a 70 mm mismatch gradually imposed. On average, participants recalibrated both vision and proprioception *(Δh*_*V*_ = 38.1 mm, *Δh*_*P*_ = 17.4 mm). **iii**. Mean distance of the five sets of 5 right-handed reaches immediately following the calibration task. *First and second sets of 5 reaches were significantly different from the pre-calibration task reaches (p_adj_ < 0.05). **B. Veridical group (N=16)**. Post-calibration reaches did not differ significantly from pre-calibration task reaches. All error bars and shaded regions represent standard error.

Within each group, paired-sample t-tests were used to compare each set of post-calibration reaches (Fig. 3iii) with the pre-calibration reach set (Fig. 3i). For the Mismatch group, the first and second set of post-calibration task reaches were significantly different from the pre-calibration set (t_15_ = 3.47, p_adj_ = 0.014; t_15_ = 2.86, p_adj_ = 0.024). The third and fourth set were not significantly different from the pre-calibration reaches (t_15_ = 1.52, p_adj_ = 0.20; t_15_ = 0.76, p_adj_ = 0.46), suggesting any effect of mismatch on reach distance did not last beyond the first two sets of 5 reaches. For the Veridical group, none of the post-calibration task sets of reaches differed significantly from the pre-calibration set (all p > 0.5). Finally, a two-sample t-test comparing the pre-calibration reaches across groups did not yield any evidence that the two groups reached different distances prior to the calibration task (t_30_ = 0.26, p = 0.80).

For the mismatch group, we compared participants’ change in right-hand reach undershoot from the last 5 pre-calibration reaches to the first 5 post-calibration reaches with total recalibration magnitude (visual plus proprioceptive) and with proprioceptive recalibration (Fig. 4). Change in reach undershoot was significantly smaller than total recalibration (t_15_ = −4.32 p_adj_ = 0.0012), but not significantly different from proprioceptive recalibration (t_15_ = 0.46, p = 0.65). This could indicate that change in reach performance does not reflect the sum of visual and proprioceptive recalibration (Fig. 1C).

**Figure 4.**
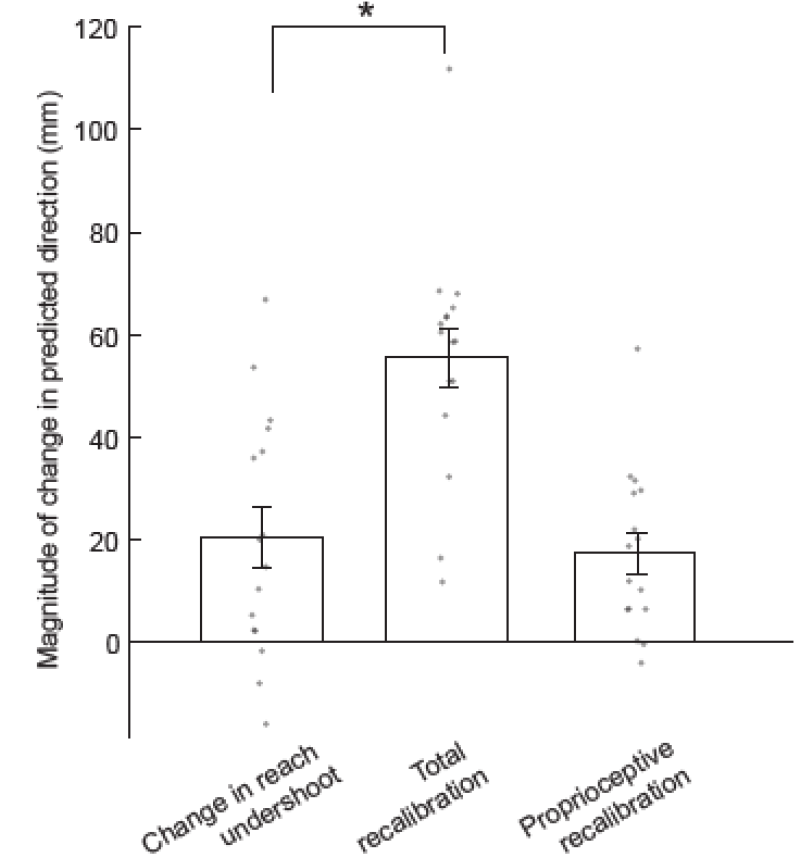
In the Mismatch group, reach undershoot changed by an average magnitude of 20.9 mm in the predicted direction. This magnitude was significantly smaller than the total magnitude of recalibration (visual plus proprioceptive), which averaged 55.5mm (*p_adj_ < 0.05). Change in reach undershoot did not differ significantly from the magnitude of proprioceptive recalibration, which averaged 17.4 mm.

Participants were questioned about their experience at the conclusion of the session. Most participants did not perceive any forward mismatch between the visual and proprioceptive targets during the calibration task. Two participants in each group reported perceiving such a mismatch, which is consistent with our previous work (Hsiao et al. 2022). Participants were also asked to rate their quality of sleep the previous night, their level of attention during the task, and how fatigued they felt after the task, on a scale of 1 to 10. We found no indication that these might differ across groups. Sleep was rated 6.9 ± 1.0 (mean ± 95% CI) by the Mismatch group and 6.6 ± 0.8 by the veridical group. Attention was rated 7.2 ± 0.5 by the Mismatch group and 7.8 ± 0.6 by the veridical group. Fatigue was rated 4.1 ± 1.4 by the Mismatch group and 3.8 ± 0.8 by the veridical group.

## 4. DISCUSSION

Here we asked whether reaching movements show evidence of change after visuo-proprioceptive recalibration in hand position estimates. The Mismatch group, after exposure to a gradual 70 mm visuo-proprioceptive mismatch, made significantly shorter reaching movements. The magnitude of change in reach distance was similar to the magnitude of proprioceptive recalibration. However, reaching performance returned to baseline levels after only 10 reaches. The Veridical group, after exposure to veridically-aligned visual and proprioceptive cues about the hand, showed no evidence of a change in reach distance. Taken together, these results suggest that visuo-proprioceptive recalibration of hand estimates does affect reaching movements, but only briefly.

### Reach distance affected by visuo-proprioceptive recalibration

We predicted that reach distance would shorten after exposure to a 70 mm forward displacement of visual cues from proprioceptive cues about the hand. We reasoned that with this direction of cue conflict, proprioceptive recalibration would be expected in the forward direction. In other words, participants would come to feel that their target hand was further forward than it actually was. They would then execute reaches of smaller magnitudes, feeling there was less distance to travel from their proprioceptively-perceived hand position to the visual target. Results of the present study support this prediction. The Mismatch group reached shorter distances after exposure to the visuo-proprioceptive mismatch, compared to baseline. Importantly, the Veridical group showed no change in reaching after exposure to veridical visuo-proprioceptive cues, indicating that the reach distance change was specific to the experience of a visuo-proprioceptive cue conflict.

An important question to consider is whether it is possible that the reaching hand experienced motor adaptation during the calibration task, or any other motor learning process that could affect reach distance; this would confound our interpretation of the role of proprioceptive recalibration in the change in reach distance. Importantly, the calibration task was designed to preclude any such confound: participants never received any information about where their indicator finger landed in relation to the target, so there was no error signal that could drive motor adaptation. In addition, participants were explicitly instructed to place their indicator finger at the perceived target location, with no time constraints. The one form of motor learning that should be possible in these conditions is for the indicator finger’s pointing movements to become less variable across the calibration task. In other words, if the participant executes a movement of their indicator finger and proprioceptive feedback from the indicator hand suggests the finger did not land in the planned position, the brain could fine-tune the motor command to make more accurate predictions. However, this form of learning should not be considered a confound, as it would occur similarly in both the Misaligned and Veridical groups.

### Implications for the sensorimotor map and other body representations

Our prediction of shortened reach distance after visuo-proprioceptive recalibration was based on the idea that perception of body parts is closely linked to motor control, and that reach planning would therefore take into account any changes in perceived hand position in order to maintain movement accuracy. Consistent with this idea, we have previously observed changes in the excitability of primary motor cortex (M1) that were specifically related to visuo-proprioceptive recalibration, after controlling for motor behavior (Munoz-Rubke et al. 2017). Furthermore, M1 changes were somatotopically focal, limited to the M1 representation of the finger that experienced visuo-proprioceptive cue conflict (Mirdamadi et al. 2022). Findings like these are suggestive of a close relationship between perception of hand position and motor execution involving that hand.

On the other hand, there is evidence to suggest that changes in perceived hand position may affect a body representation that functions separately from the body representation used to control movement. Paillard (1999) refers to these representations as the body image and the body schema, respectively, based on a double dissociation observed in neuropsychological patients (Paillard 1999). In other words, some patients can correctly use information about their body positioning to move (intact body schema), but do not correctly perceive their body positioning (disrupted body image), while others have the opposite problem (Paillard 1999). This allows us to infer that there are at least two dissociable body representations, although some have suggested that body image should be further divided (Kammers et al. 2009; de Vignemont 2010).

The literature includes examples where a perceptual manipulation clearly does not affects motor performance (Kammers et al. 2009), examples where it clearly does (Cressman and Henriques 2010; Salomonczyk et al. 2013), and also gray areas (Botvinick and Cohen 1998). Some of this literature uses the Rubber Hand Illusion (RHI), in which synchronous stroking of a seen fake arm and the felt real arm creates the illusion of body ownership over the fake arm (Botvinick and Cohen 1998). Kammers et al. (2009) found that perceptual bodily judgments were sensitive to the RHI, but ballistic motor responses were not. The authors thus concluded that the illusion affected the body image, but not the body schema (Kammers et al. 2009). However, while the RHI entails a visuo-proprioceptive discrepancy like the present study, there are also important differences to be considered. Our paradigm lacks synchronous tactile stimulation, and the visual stimulus is reduced to a disembodied white disc. The RHI is associated with “visual capture”, where the visual signal is so much stronger than the proprioceptive signal that most recalibration is likely proprioceptive rather than visual. In contrast, our paradigm is associated with a slightly stronger weight of proprioception compared to vision, and greater visual recalibration than proprioceptive (Block and Sexton 2020; Liu et al. 2018). Indeed, Kammers et al. (2009) suggested that weighting of vision vs. proprioception in multisensory integration could explain their results, with perceptual judgments relying heavily on vision and ballistic movements relying on proprioception.

Even within the RHI literature, there are grey areas in terms of which body representation appears affected. Kammers et al. (2009) noted that the classic RHI study by Botvinick and Cohen (1998) assessed illusion strength by asking participants to make pointing movements to where they perceived their other hand. The pointing movements were clearly sensitive to the illusion (Botvinick and Cohen 1998), unlike the ballistic movements used by Kammers et al. (2009). However, the pointing movements may have accessed a perceptual judgment in a way that ballistic movements do not (Kammers et al. 2009).

On the other end of the spectrum, there is evidence that exposure to a visuo-proprioceptive mismatch in a cursor rotation task robustly affects reaching movement (Cressman and Henriques 2010; Salomonczyk et al. 2013). The cursor rotation paradigm is often used to elicit visuomotor adaptation: participants move to a visual target while the corresponding cursor deviates by some angular magnitude. This results in systematic movement errors, which are reduced by trial-and-error adaptation of the motor command. Proprioceptive recalibration also occurs, due to both sensory prediction errors and the cross-sensory mismatch created by the deviation of the cursor from true hand position (Rossi et al. 2021). Cressman and Henriques (2010) modified the cursor rotation paradigm to remove the movement errors that could drive visuomotor adaptation; instead, participants moved their hand actively or passively along a set channel that was gradually deviated from the cursor, which always went straight to the target. After this exposure, participants made self-guided reaches with no cursor. These no-cursor reaches showed a directional change after exposure to the visuo-proprioceptive mismatch, and the change in reach direction was correlated with participants’ magnitude of proprioceptive recalibration (Cressman and Henriques 2010).

While a visuo-proprioceptive mismatch created in the context of cursor rotation seems to clearly alter the body schema, we must be cautious in comparing cursor rotation findings with the present study. The nature of a cursor rotation is that it exists in the context of target-directed movement: there is no visuo-proprioceptive mismatch at the home position, and the mismatch linearly increases as the person approaches the target. In addition, the mismatch is applied in body-centered coordinates; moving the hand to the right would result in a mismatch of the opposite direction in extrinsic space compared to moving the hand to the left. In contrast, the present study imposed the mismatch with the hand positioned in a static location, and that hand made no movements toward a target while the mismatch was imposed.

In sum, the present paradigm has features in common with both the RHI and the cursor rotation task, but also features that differ. Based on the above studies, we would suggest that our visuo-proprioceptive mismatch task affected primarily the body image, as participants were explicitly asked to indicate their perceived positions of static hand targets as accurately as possible without regard to speed (Botvinick and Cohen 1998; Kammers et al. 2009). The straight-ahead reaches presumably accessed the body schema, which makes it surprising that we observed a shortening of reaches.

However, it should be noted that the body schema and body image likely interact with each other (de Vignemont 2010), making it challenging to draw more specific conclusions.

### Magnitude of reach distance reduction was similar to proprioceptive recalibration magnitude, but transient

We hypothesized that in the absence of visual feedback about the hand during reaching, only the proprioceptive recalibration would contribute to target undershoot. Results were consistent with this hypothesis. However, we also considered the possibility that recalibration of visual estimates of the hand could include everything in the visual scene. In other words, it is possible that when visual recalibration occurs, people interpret that as the visual scene being closer than it looks, rather than specifically visual information about the hand. However, if that were the case, we would expect change in reach undershoot to be larger than proprioceptive recalibration, and more similar to total recalibration.

It is interesting to note that the reduction in reach distance was no longer detectable after only 10 reaches. Motor adaptation, in contrast, can be retained even after a year (Yamamoto et al. 2006). Proprioceptive recalibration that results from motor adaptation can, itself, still be evident after 24 hours (Maksimovic and Cressman 2018; Nourouzpour et al. 2015). There are several possible interpretations of this difference. First, more exposure to a visuo-proprioceptive mismatch might result in longer-lasting reduction in reach distance; there were only 42 visuo-proprioceptive exposure trials in the present study, while motor adaptation studies frequently involve hundreds of trials. Second, perhaps the active movement stimulates proprioceptors in a way that overrides proprioceptive recalibration generated at a static position. This possibility could be considered consistent with RHI studies that found the illusion is reduced by active movement of the stimulated hand (Kammers et al. 2009). Third, perhaps proprioceptive recalibration, induced with only a cue conflict and not motor adaptation, is itself transient, so effects on reach distance are also transient. However, evidence suggests that proprioceptive recalibration induced with a cue conflict is robustly retained 24 hours later (Wali et al. 2022). Finally, if proprioceptive recalibration lasts longer than 10 trials, it is possible the effect on reach distance does not. This would be consistent with the involvement of multiple body representations: proprioceptive recalibration might occur in the body image, which only briefly interacts with the body schema used to plan reaches. Indeed, different body representations have been associated with different dynamics and timescales (de Vignemont 2010). Of course, these possible interpretations are not mutually exclusive. Further studies are needed to better understand what factors influence the time course of visuo-proprioceptive recalibration effects on movement.

## Conclusions

We predicted that straight-ahead reach distance would shorten after exposure to a forward displacement of visual cues from proprioceptive cues about the hand, which leads to proprioceptive recalibration in the forward direction. Results support this prediction, but the reduction of reach distance was transient. This is consistent with some degree of separation between body representations for perception and action.

## CRediT author statement

HJB: Conceptualization, Methodology, Software, Formal analysis, Writing – Review &Editing, Visualization, Supervision, Funding acquisition.

YL: Software, Formal analysis, Investigation, Writing – Original Draft, Visualization.

